# Codon language model scores provide information beyond protein language models for missense variant interpretation

**DOI:** 10.1101/2025.03.12.642937

**Authors:** Ruyi Chen, Nathan Palpant, Gabriel Foley, Mikael Bodén

## Abstract

Predicting variant effects remains a central challenge in genomics. Protein language models (PLMs) capture amino-acid-level sequence constraints, whereas codon language models operate on coding sequences and may retain information that is lost upon translation. Here, we tested whether scores from the codon language model CaLM provide predictive information beyond protein-level representations for missense-variant interpretation. Across 71,436 ClinVar missense variants from 11,554 genes, adding CaLM to PLM baselines produced modest but reproducible improvements under gene-held-out cross-validation. PLM-only ensemble controls and explicit mutational-context analyses indicated that this improvement could not be explained solely by generic ensembling or simple codon-substitution features. Aggregating CaLM probabilities across synonymous codons attenuated codon-degeneracy-associated discordance while preserving most of the broader differences between CaLM and PLM scores. Gene-level analyses further showed that CaLM contribution varied continuously across genes and depended partly on the protein-model background. Across ClinMAVE functional assays, however, improvements were less consistent, indicating that codon–protein complementarity is context dependent rather than universal. Together, these results identify a modest but reproducible component of variant-effect information in CaLM-derived codon-level scores that is not fully captured by protein-level language-model representations.

## 1 Introduction

Interpreting the functional consequences of genetic variants is a central goal in genomics, yet remains challenging at scale. Deep learning models increasingly enable zero-shot estimation of variant effects by extracting evolutionary constraints directly from biological sequences [1–3]. In many sequence language models, variant effects can be estimated from the change in probability assigned to the mutant relative to the wildtype (wt) state in its native sequence context. This difference is commonly expressed as a log-likelihood ratio (LLR), yielding scores that correlate with experimental fitness and clinical pathogenicity [1, 4].

Most sequence-based approaches to missense-variant interpretation operate at the protein level or rely primarily on protein-centred representations. These include protein language models (PLMs) trained on individual protein sequences [1, 3, 5], multiple-sequence-alignment-based models [2, 6], and structure-informed models [7– 10]. Such approaches have been highly successful because many pathogenic missense variants alter protein stability, folding, or function. Coding sequences, however, are not interchangeable encodings of proteins. Even when amino-acid identity is unchanged, synonymous codons differ in nucleotide composition and local coding context, while the same amino-acid substitution can arise through distinct codon-level changes. Synonymous codon choice, local nucleotide context, and genetic-code degeneracy have been associated with mutation, translation, and gene regulation [11, 12], yet these properties are collapsed when coding sequences are represented solely at amino-acid resolution.

Codon language models (CLMs) instead model coding sequences in their native triplet representation and have achieved performance comparable to PLMs on several protein-property prediction tasks [13]. Comparable performance, however, does not establish that CLMs capture codon-level information that is specifically relevant to variant interpretation. Some signals learned by codon models have been attributed to codon-usage patterns that reflect species of origin rather than finer functional constraints [14]. It therefore remains unclear whether codon-level models capture information relevant to missense-variant effects and, more importantly, whether such information remains informative once strong protein-level predictors are taken into account. Conceptually, codon- and protein-level models describe closely related sequence modalities that share substantial biological information while retaining distinct local constraints (Figs. 1A and B). This motivates a central question: do codon-level likelihoods provide pathogenicity information beyond that already captured by PLMs, or do they primarily recapitulate the same protein-level signal?

**Fig. 1.**
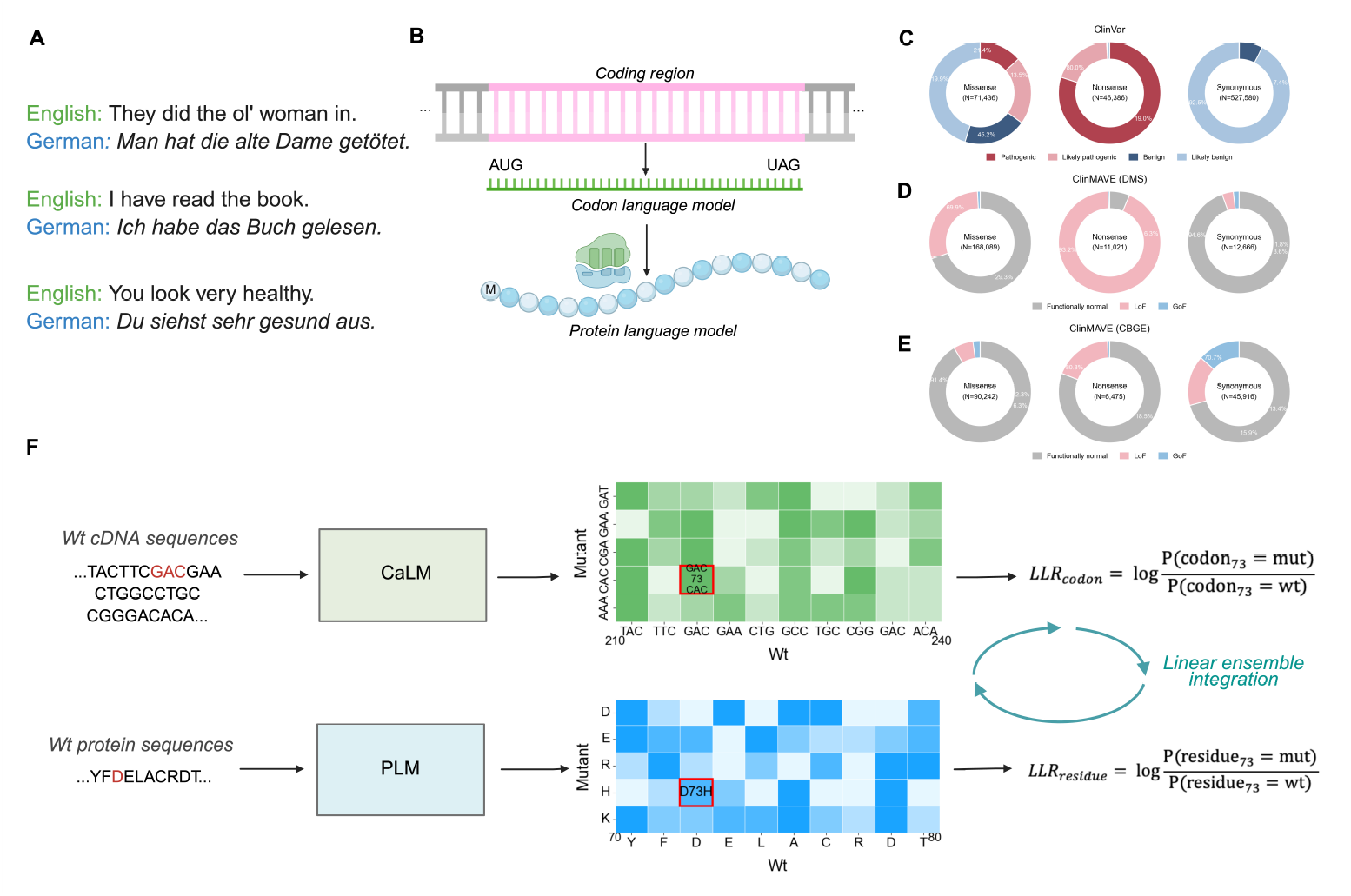
Codon- and protein-level models capture partially distinct constraints across coding sequences and their protein products. **(A)** Natural-language analogy for cross-modal representation. English and German sentences can convey the same core meaning while differing in syntax and local expression. **(B)** Schematic analogy for codon- and protein-level sequence models. CaLM scores coding sequences in codon space, whereas PLMs score the translated protein sequence in amino-acid space; these representations preserve overlapping but non-identical information. **(C–E)** Dataset composition and label distributions for ClinVar and ClinMAVE. Panels show the distribution of variant classes and ground-truth labels in (C) ClinVar, (D) ClinMAVE DMS, and (E) ClinMAVE CBGE datasets. **(F)** Ensemble-evaluation framework. CaLM processes wt cDNA sequences to compute codon-level effect scores, while PLMs process protein sequences to compute residue-level effect scores. Scores are evaluated individually and integrated using cross-validated convex linear ensembles to predict variant pathogenicity or functional labels. This figure was created with BioRender.com.

Here, we systematically tested whether CaLM-derived codon-level LLRs provide complementary information for missense-variant interpretation beyond amino-acid-level PLM scores. Using ClinVar missense variants [15], we compared CaLM with three PLMs, ESM-2 (150M), ESM-2 (650M), and ESM-1b (650M), both individually and in controlled multimodal ensembles. Same-modality PLM controls were used to distinguish cross-modal complementarity from the generic improvement expected when combining multiple predictors. We then tested whether the additional CaLM signal could be explained by explicit mutational-context features or by information retained after collapsing codon probabilities into amino-acid space. We further examined heterogeneity in CaLM contribution across genes and PLM backgrounds. Finally, using ClinMAVE datasets derived from deep mutational scanning (DMS) and CRISPR-based genome editing (CBGE) [16], we assessed whether the inferred contribution of codon-level information varied across functional classes and experimental contexts. Together, these analyses show that CaLM-derived codon-level likelihoods contain a modest but reproducible component of variant-effect information that is not fully captured by protein-level language-model scores.

## 2 Results

### 2.1 Codon-level scores provide partially non-redundant information beyond PLMs

To test whether CaLM-derived codon-level scores provide information beyond PLM-derived amino-acid scores, we collected 71,436 ClinVar missense variants, comprising 24,933 pathogenic and 46,503 benign variants across 11,554 genes (Fig. 1C and Supplementary Table S1). LLR-derived pathogenicity scores were evaluated individually and in convex linear ensembles using 10-fold gene-held-out cross-validation, with all ensemble optimisation restricted to the training genes (Fig. 1F). Model comparisons were based on the pooled out-of-fold difference in the area under the receiver operating characteristic curve (ΔAUROC), with 95% confidence intervals (CIs) estimated by gene-cluster bootstrap (see “Methods – Model evaluation and ensemble optimisation”).

The pooled out-of-fold AUROCs were 0.836 for ESM-2 (150M), 0.893 for ESM-2 (650M), 0.927 for ESM-1b (650M), and 0.840 for CaLM (Fig. 2A). Among PLM-only combinations, adding ESM-2 (150M) to ESM-2 (650M) produced only a small improvement over ESM-2 (650M) (ΔAUROC = 0.0007; 95% CI, 0.00004–0.00129). Combining ESM-2 (650M) with ESM-1b (650M) produced a substantially larger gain relative to ESM-2 (650M) (ΔAUROC = 0.0415; 95% CI, 0.0350–0.0481), although the improvement relative to the stronger constituent, ESM-1b (650M), was 0.0075 (95% CI, 0.0054–0.0098). CaLM provided additional predictive value across multiple PLM backgrounds. Adding CaLM to ESM-2 (650M) increased AUROC by 0.0183 (95% CI, 0.0149–0.0220), while adding CaLM to ESM-1b (650M) increased AUROC by 0.0060 (95% CI, 0.0049–0.0073). CaLM also improved the combined ESM-2 (650M) and ESM-1b (650M) baseline by 0.0031 (95% CI, 0.0022–0.0040; Fig. 2A). Thus, the magnitude of the CaLM gain decreased as the protein baseline strengthened, but remained positive even when added to the strongest PLM-only ensemble. The direction of improvement was also consistent across cross-validation folds (Supplementary Fig. S1), and variant-wise cross-validation reproduced the same qualitative model ordering (Supplementary Fig. S2).

**Fig. 2.**
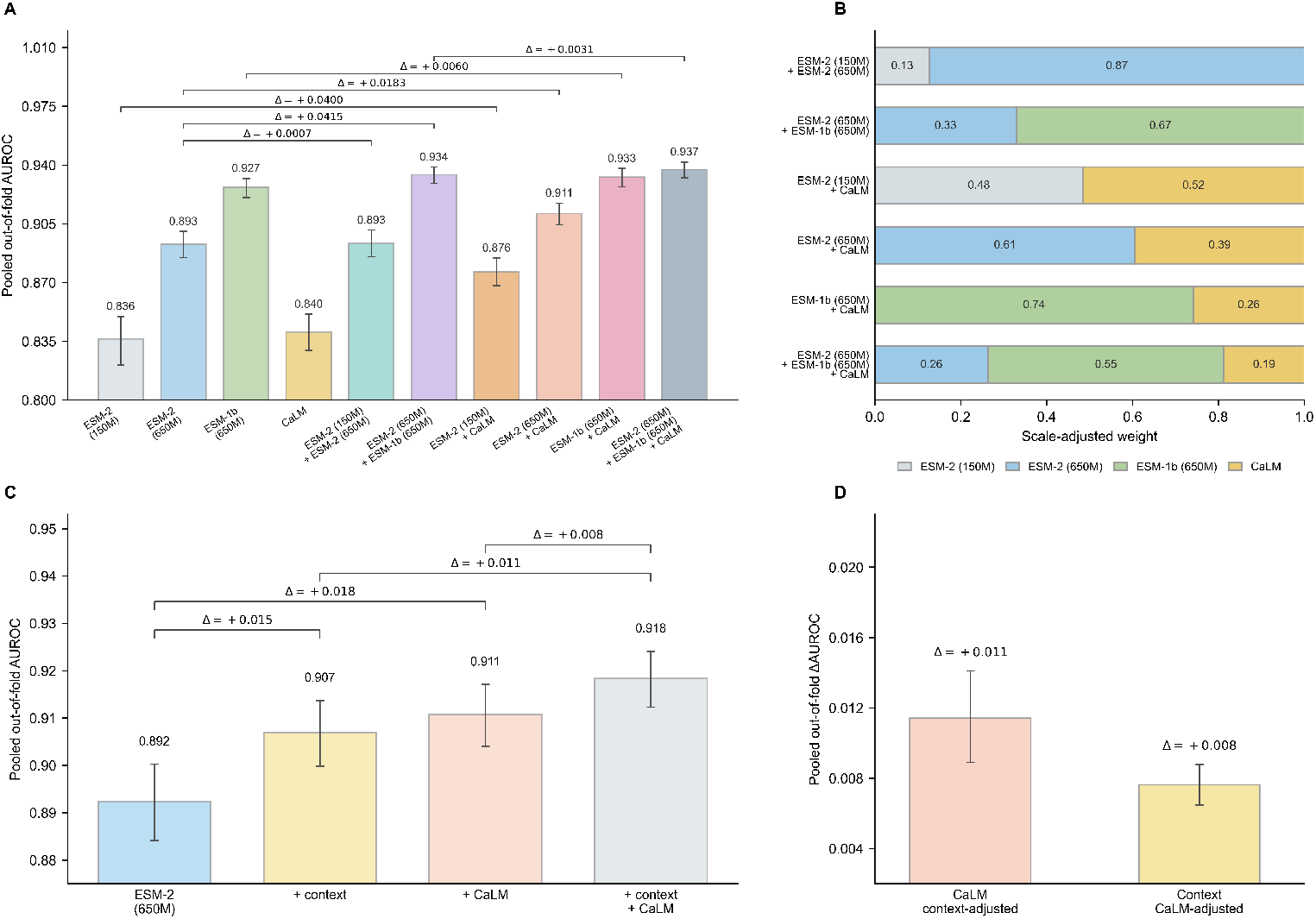
CaLM provides complementary predictive information beyond PLM scores and explicit mutational-context features. **(A)** Pooled out-of-fold AUROC for individual models and model combinations under 10-fold gene-held-out cross-validation. Error bars indicate 95% gene-cluster bootstrap CIs. **(B)** Scale-adjusted ensemble weights for the model combinations shown in panel A. Stacked bars show the mean scale-adjusted coefficient assigned to each model across the 10 cross-validation folds. **(C)** Pooled out-of-fold AUROC for ESM-2 (650M) alone and after adding mutational-context features, CaLM, or both. Error bars indicate 95% gene-cluster bootstrap CIs. **(D)** Conditional gains in pooled out-of-fold AUROC after accounting for the alternative information source. Points show ΔAUROC values, and error bars indicate 95% gene-cluster bootstrap CIs.

We next used PLM-only ensembles as same-modality placeholder controls to ask whether the incremental benefit of CaLM exceeded that obtained by adding another protein-level scorer. Relative to ESM-2 (650M), adding CaLM produced a larger gain (ΔAUROC = 0.0183) than adding ESM-2 (150M) (ΔAUROC = 0.0007). The CaLM gain was also larger than the incremental improvement of the ESM-2 (650M) and ESM-1b (650M) ensemble over its stronger constituent, ESM-1b (650M) (ΔAUROC = 0.0075). This distinction diminished, however, when ESM-1b (650M) itself served as the baseline: adding CaLM increased AUROC by 0.0060, comparable in magnitude to the 0.0075 gain obtained by adding ESM-2 (650M). Thus, the apparent advantage of cross-modal ensembling depended on baseline strength rather than uniformly exceeding that of PLM-only combinations. CaLM nevertheless contributed despite having substantially lower individual AUROC than ESM-1b (650M). Its scores were also less correlated with ESM-2 (650M) than those of either added PLM (Supplementary Table S4), consistent with CaLM capturing a partially distinct component of the predictive signal. Fold-level sensitivity analyses showed the same qualitative pattern (Supplementary Fig. S3).

The scale-adjusted ensemble weights were consistent with the AUROC-based comparisons. In PLM-only ensembles, larger weights were assigned to the stronger protein model, whereas ensembles containing CaLM retained a non-zero weight on the codon-level score across PLM backgrounds (Fig. 2B and Supplementary Fig. S4). The CaLM coefficient decreased as stronger protein-level information was introduced, consistent with partial overlap rather than complete independence between the two modalities.

Together, these results indicate that CaLM-derived codon-level scores contain a modest but reproducible component of predictive information that is not fully captured by PLM scores. Same-modality placeholder controls further suggest that this improvement is not solely attributable to the generic benefit of adding another predictor, although the distinction from the strongest PLM placeholder becomes small when the protein baseline is already strong.

### 2.2 CaLM provides predictive information beyond explicit mutational-context features

We next tested whether the incremental CaLM signal could be explained by explicitly encoded properties of the underlying codon substitution. Using ESM-2 (650M) as a fixed protein-model background, we evaluated four feature configurations on the same ClinVar missense dataset under 10-fold gene-held-out cross-validation: ESM-2 (650M) alone, ESM-2 (650M) with mutational-context features, ESM-2 (650M) with CaLM, and ESM-2 (650M) with both mutational-context features and CaLM. All configurations were evaluated within the same L2-regularised logistic-regression framework. The mutational-context covariates and methodological details are described in “Methods – Codon-level mutational-context control analysis” and Supplementary Table S5.

Within this logistic-regression control framework, ESM-2 (650M) alone achieved a pooled out-of-fold AUROC of 0.893. Adding mutational-context features increased AUROC to 0.907 (ΔAUROC = 0.0146), whereas adding CaLM increased it to 0.911 (ΔAUROC = 0.0183; Fig. 2C). Including both sources produced the highest AUROC, 0.918 (ΔAUROC = 0.0260). The combined improvement was smaller than the sum of the two individual gains, consistent with partial overlap between the information captured by CaLM and the explicit mutational-context features, although AUROC gains are not expected to combine additively.

We next quantified the conditional contribution of each source after accounting for the other. Adding CaLM to the ESM-2 (650M) plus mutational-context model increased AUROC by 0.0114 (95% CI, 0.0089–0.0141). Conversely, adding mutational-context features to the ESM-2 (650M) plus CaLM model increased AUROC by 0.0076 (95% CI, 0.0065–0.0088; Fig. 2D). The direction of both conditional improvements was also consistent across all 10 held-out folds (Supplementary Fig. S5).

These results indicate that CaLM scores share predictive information with explicit codon-substitution features but are not fully accounted for by them. Together with the PLM-combination analysis, these controls support a modest but partially distinct contribution of CaLM-derived codon-level scores beyond both PLM-derived scores and the measured mutational context.

### 2.3 CaLM contribution varies across functional assays

We next tested whether CaLM-derived codon-level signals generalized beyond Clin-Var across functional classes, experimental platforms, and PLM backgrounds. Using ClinMAVE missense variants from DMS and CBGE experiments [16], we evaluated functionally normal versus loss-of-function (LoF) and gain-of-function (GoF) variants under gene-held-out cross-validation (Figs. 1D and E; Supplementary Table S2). CaLM was combined separately with ESM-2 (650M) and ESM-1b (650M) using the same convex ensemble procedure. Ensemble weights were scale-adjusted within training folds for cross-context comparison, while performance was evaluated from the original pooled out-of-fold predictions with uncertainty estimated by gene-cluster bootstrap (Supplementary Table S3).

For LoF discrimination, both PLMs achieved higher AUROC than CaLM when evaluated individually. In DMS, adding CaLM produced positive gains with both ESM-2 (650M) (ΔAUROC = 0.0101; 95% CI, 0.0030–0.0172) and ESM-1b (650M) (ΔAUROC = 0.0161; 95% CI, 0.0069–0.0263; Fig. 3B). In CBGE, the corresponding CIs included zero, providing no robust evidence of improved held-out performance. The scale-adjusted CaLM weights also varied across platforms and PLM backgrounds (Fig. 3A), illustrating that a non-zero training-optimised weight does not necessarily translate into a generalisable gain on held-out genes.

**Fig. 3.**
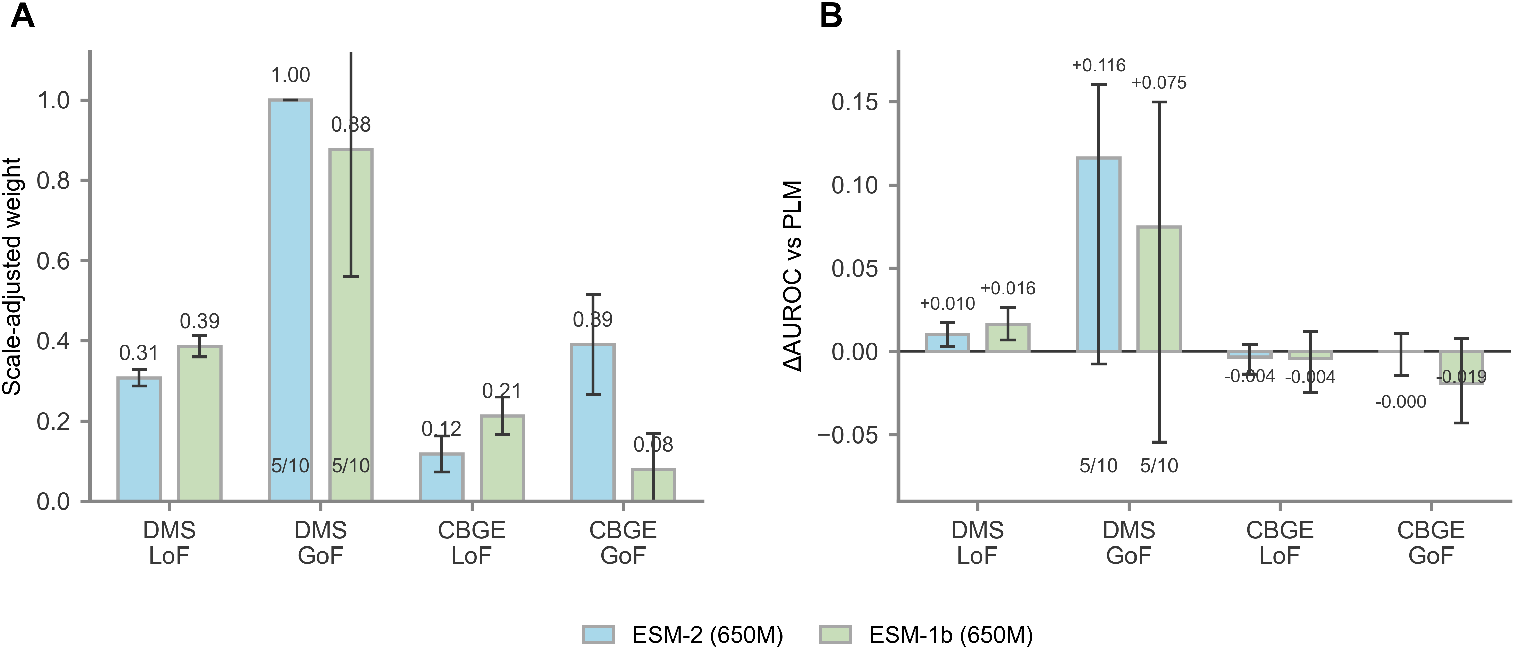
CaLM contribution varies across functional classes, assay platforms, and protein-model backgrounds. **(A)** Scale-adjusted CaLM weights for functionally normal versus LoF and functionally normal versus GoF comparisons in the DMS and CBGE datasets. CaLM was combined with either ESM-2 (650M) or ESM-1b (650M) using convex linear ensembles. Bars show means across cross-validation folds, and error bars indicate the standard deviation across folds. **(B)** Corresponding pooled out-of-fold AUROC gain of the CaLM–PLM ensemble relative to the PLM alone. Error bars indicate 95% gene-cluster bootstrap confidence intervals, and the horizontal line denotes zero gain. The “5/10” labels indicate that five of the ten folds contained both classes for the fold-level analysis.

GoF comparisons were substantially less stable. In DMS, only 5 of the 10 held-out folds contained both classes and therefore permitted fold-level AUROC calculation. The pooled out-of-fold gains from adding CaLM were positive with ESM-2 (650M) (ΔAUROC = 0.1161) and ESM-1b (650M) (ΔAUROC = 0.0747), but the corresponding gene-bootstrap confidence intervals included zero (Fig. 3B). The optimised CaLM weight in the ESM-2 (650M) background reached the boundary value of 1.00 in all training folds, whereas the CaLM weights in the ESM-1b (650M) background were more variable. Given the sparse class representation and the lack of robust held-out improvement, these high weights are better interpreted as training-set preference than as evidence that codon-level scores dominate GoF prediction.

For CBGE-GoF, adding CaLM likewise did not produce a robust improvement with either PLM background, with both gene-bootstrap confidence intervals including zero. Fold-level GoF gains and optimised weight distributions are shown in Supplementary Figs. S6 and S7. Together, these analyses further illustrate that optimised ensemble weights alone do not establish a generalisable contribution on held-out genes.

Across ClinMAVE functional assays, the inferred contribution of CaLM depended on functional class, experimental platform, and PLM background. CaLM provided modest but reproducible gains in some LoF settings, whereas large optimised weights in the sparser GoF analyses did not translate into robust held-out improvements. Thus, the complementary contribution observed in ClinVar was not uniformly reproduced across functional-assay settings, indicating that the predictive value of CaLM-derived codon-level scores is context-dependent and less stable in ClinMAVE.

### 2.4 Codon-level scores distinguish nonsense from synonymous variants

Residue-level PLM LLRs are defined for amino-acid substitutions, leaving synonymous variants without an amino-acid change to score, while nonsense variants fall outside the missense-scoring framework used here. We therefore examined whether CaLM provided a meaningful scoring axis for variant classes not directly represented by these residue-level PLM scores. Across ClinVar, ClinMAVE-DMS, and ClinMAVE-CBGE, CaLM assigned substantially lower LLRs to nonsense than to synonymous variants (two-sided Mann–Whitney U tests; *p <* 10^−300^). Discriminating functional effects within either class was considerably more difficult. LLR distributions for pathogenic and benign synonymous variants overlapped substantially in ClinVar (Fig. 4A), while separation of functionally normal variants from LoF or GoF variants was limited in both DMS and CBGE (Fig. 4B). Nevertheless, the broad separation between non-sense and synonymous variants was consistent across datasets, suggesting that the distinction was not specific to a particular assay or annotation source.

**Fig. 4.**
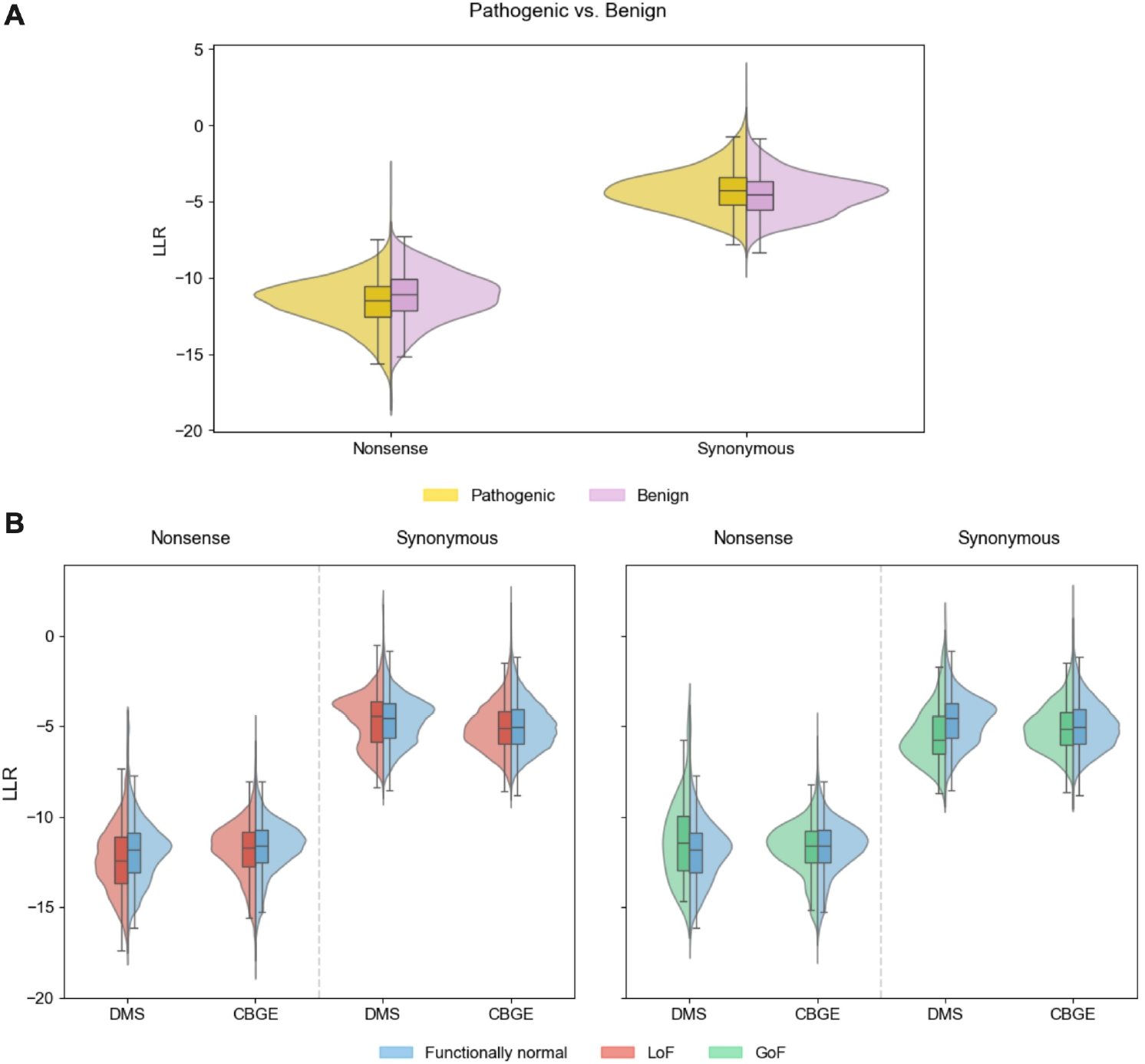
Codon-level scores distinguish nonsense from synonymous variants. **(A)** CaLM LLR distributions for ClinVar nonsense and synonymous variants, stratified by clinical classification as pathogenic or benign. **(B)** CaLM LLR distributions for ClinMAVE nonsense and synonymous variants across functional categories and experimental platforms. The left sub-panel compares functionally normal and LoF variants, whereas the right sub-panel compares functionally normal and GoF variants. Within each sub-panel, variants are stratified by sequence consequence and grouped by DMS or CBGE platform. Split violin plots show the full score distributions, with internal box plots indicating the median and interquartile range.

This category-level separation is biologically expected: premature stop codons are strongly disfavoured within coding sequences, whereas many synonymous substitutions preserve the encoded protein sequence and are comparatively tolerated. We therefore interpret this result primarily as a face-validity observation rather than evidence that CaLM provides fine-grained functional prediction within these variant classes. CaLM nevertheless supplies a codon-level scoring axis that is unavailable to the residue-level PLM framework used here, while its ability to resolve functional heterogeneity among synonymous or nonsense variants remains limited.

### 2.5 Synonymous-codon probability splitting contributes to cross-modal discordance

To examine one potential source of CaLM–PLM disagreement, we asked whether discordance arose partly because CaLM distributes probability across synonymous codons encoding the same amino acid. Using ESM-2 (650M) as the fixed PLM reference, we summed CaLM probabilities across synonymous codons and recalculated CaLM LLRs in amino-acid space. Cross-modal discordance was defined as the difference between the ESM-2 LLR and the corresponding CaLM LLR (see “Methods – Characterisation of cross-modal substitution discordance”).

We first compared discordance before and after amino-acid aggregation to assess its dependence on codon resolution. Discordance remained highly correlated across representations (Pearson *r* = 0.95; Fig. 5A), indicating that aggregation largely preserved variant rankings. However, its magnitude decreased for many variants, suggesting that synonymous-codon probability splitting accounts for part of the CaLM–ESM-2 difference. Aggregation had little effect on overall ClinVar performance, with CaLM AUROC decreasing only from 0.840 to 0.836.

**Fig. 5.**
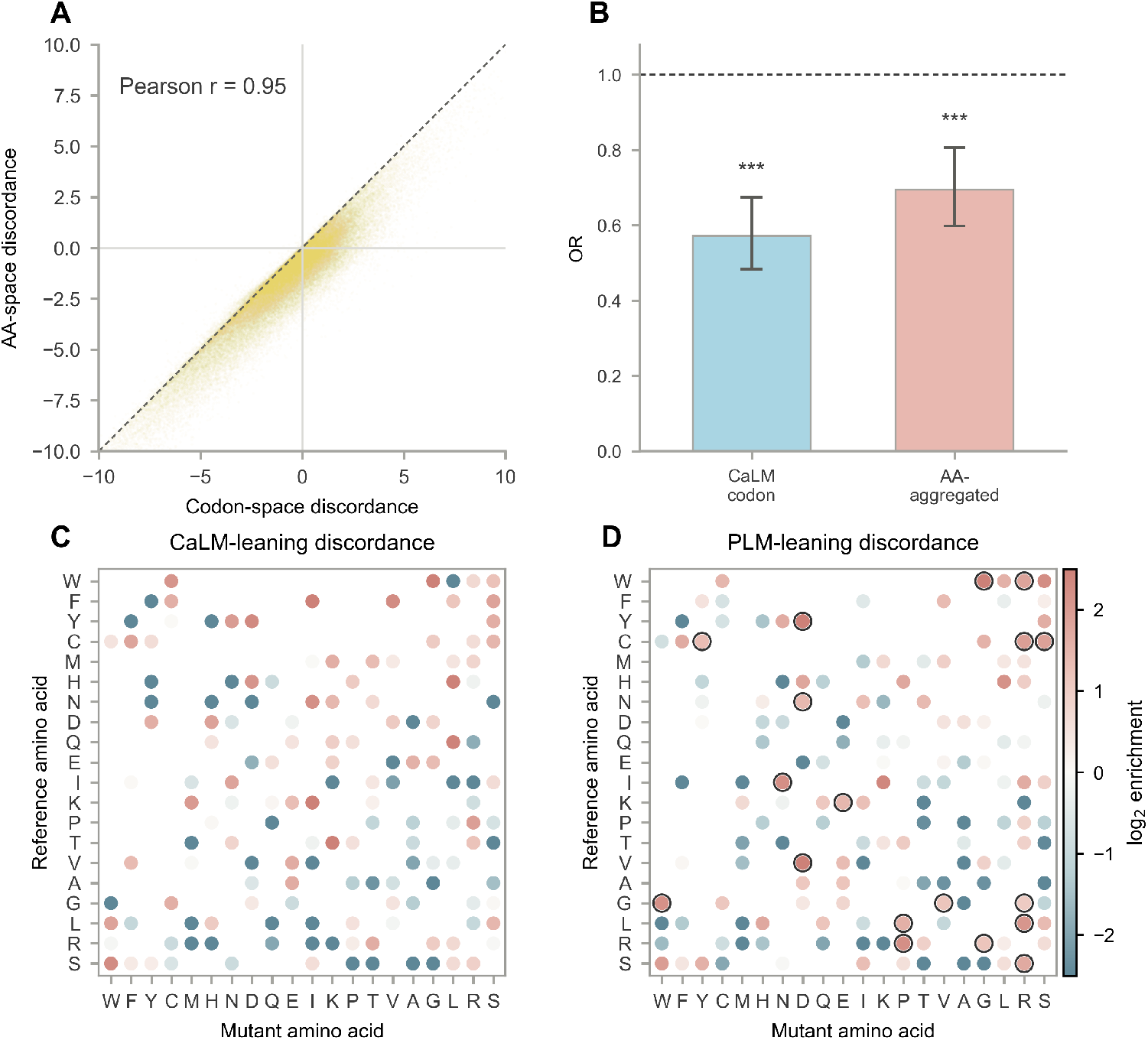
CaLM–PLM substitution discordance before and after amino-acid aggregation. **(A)** Cross-modal discordance before and after aggregating CaLM codon probabilities into amino-acid space for ClinVar missense variants. The dashed line denotes identity; points below the line indicate lower discordance after aggregation. **(B)** Association between the log reference-to-mutant codon-degeneracy ratio and CaLM-leaning discordance. Bars show odds ratios (ORs) relative to all other scored variants, with 95% CIs from gene-clustered standard errors. The dashed line denotes OR = 1; asterisks indicate *p <* 0.001. **(C–D)** Amino-acid substitution enrichment among **(C)** CaLM-leaning and **(D)** PLM-leaning variants after aggregation. Circle colour shows log_2_ enrichment relative to the full scored ClinVar missense cohort, with red indicating enrichment and blue depletion. Black outlines indicate significant enrichment after Benjamini–Hochberg correction (adjusted *p <* 0.01).

We next examined whether residual discordance after aggregation was associated with particular substitution properties. Focusing on the extreme 1% tails of the aggregated discordance distribution, 412 variants were classified as CaLM-leaning and 1,018 as PLM-leaning based on ClinVar labels. Substitutions from more to less codon-degenerate amino acids were less likely to show CaLM-leaning discordance. This association weakened after amino-acid aggregation, with the odds ratio increasing from 0.57 to 0.69 (Fig. 5B), indicating that synonymous-codon probability splitting strengthened, but did not fully explain, the relationship between codon degeneracy and cross-modal discordance.

We then asked whether the remaining discordance was concentrated in specific amino-acid substitutions. After FDR correction, no CaLM-leaning substitution was significantly enriched, whereas 18 PLM-leaning substitutions showed significant enrichment (Figs. 5C and D). Thus, residual PLM-leaning discordance was concentrated in particular substitution classes, whereas CaLM-leaning discordance was more broadly distributed. The larger number of PLM-leaning variants may also have increased power to detect enrichment.

Overall, amino-acid aggregation attenuated codon-related effects without eliminating CaLM–PLM disagreement. Synonymous-codon probability splitting therefore explains part of the observed substitution-level discordance, while additional differences persist when both models are compared at the same amino-acid resolution.

### 2.6 Gene-level codon contribution varies across genes and model backgrounds

We next examined whether the predictive contribution of CaLM-derived scores varied across genes. Using ESM-2 (650M) as the primary PLM background, we analysed 466 genes with at least five pathogenic and five benign ClinVar missense variants. Gene-level contribution was summarised by the mean scale-adjusted CaLM weight, 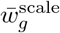, and the held-out cross-modal gain, ΔAUROC_+CaLM,*g*_ (see “Methods – Gene-level codon contribution analysis”).

Higher mean scale-adjusted CaLM weights were associated with larger held-out gains (Fig. 6A), and this relationship persisted after accounting for variant count and class balance. We therefore tested whether this association reflected CaLM-specific performance rather than differences in the PLM baseline. After adjusting for ESM-2 (650M) AUROC, the association between CaLM performance and cross-modal gain remained positive (*β* = 0.107, *p* = 2.3 *×* 10^−10^; Fig. 6B). Thus, among genes with comparable PLM performance, genes in which CaLM performed better tended to derive greater benefit from adding CaLM.

**Fig. 6.**
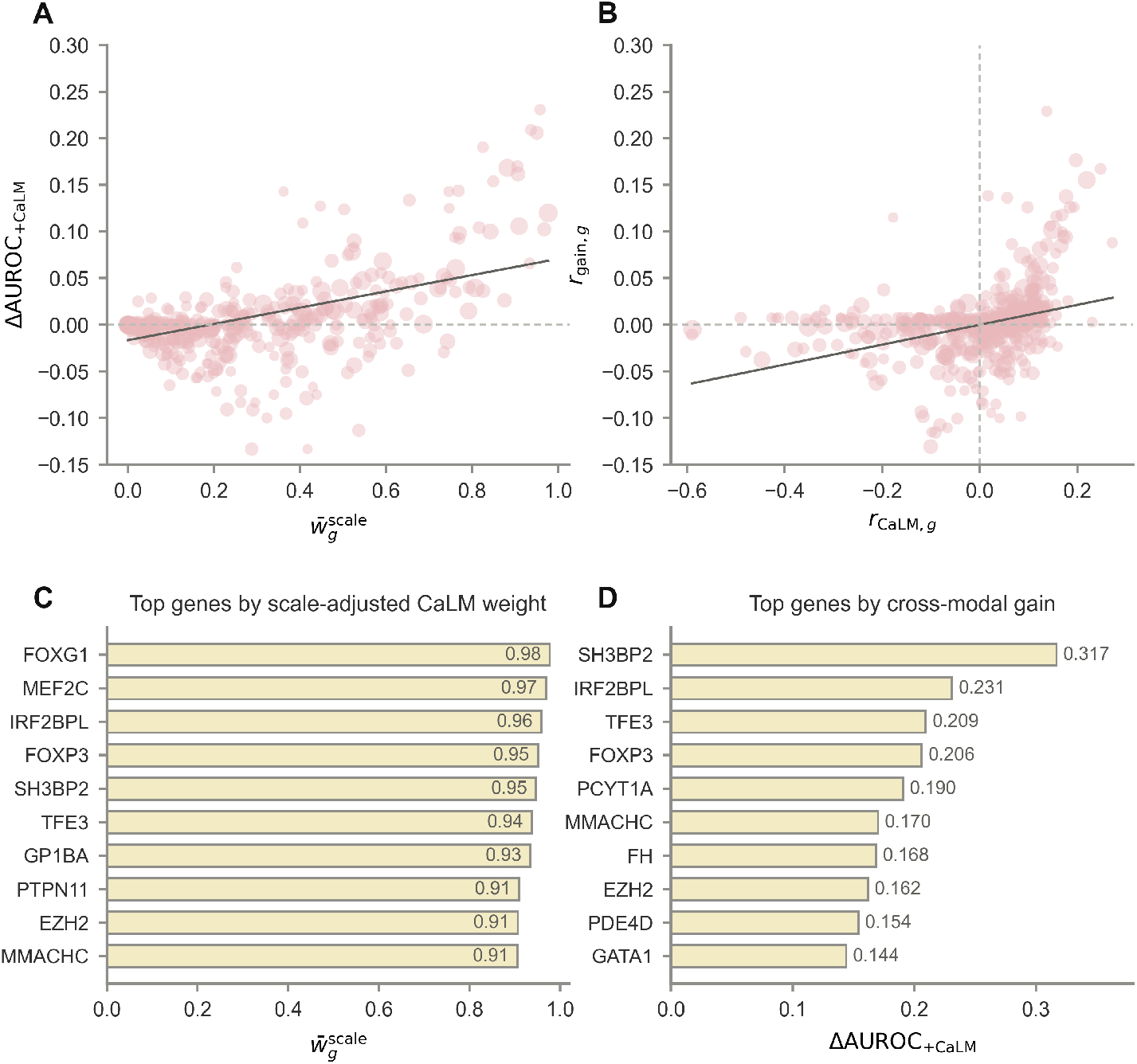
Gene-level variation in CaLM weighting and cross-modal performance gain. **(A)** Association between the mean scale-adjusted CaLM weight, 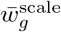, and the held-out cross-modal gain, ΔAUROC_+CaLM,*g*_ (ordinary least-squares regression: *β* = 0.087, *p* = 3.6 *×* 10^−13^, *R*^2^ = 0.23). **(B)** Partial association between CaLM performance and held-out cross-modal gain after controlling for ESM-2 (650M) AUROC. The axes show the corresponding regression residuals, *r*_CaLM,*g*_ and *r*_gain,*g*_. The partial regression coefficient was *β* = 0.107 (95% CI, 0.074–0.140; *p* = 2.3 *×* 10^−10^; partial *R*^2^ = 0.10). In **(A–B)**, each point represents one gene, point size reflects variant count, and solid lines show ordinary least-squares fits. **(C–D)** Top 10 genes ranked by **(C)** mean scale-adjusted CaLM weight and **(D)** held-out cross-modal gain.

The highest-ranked genes by mean scale-adjusted CaLM weight and held-out gain are shown in Figs. 6C and D, respectively. Several genes appeared near the top of both rankings, including *IRF2BPL, TFE3, FOXP3, SH3BP2, EZH2*, and *MMACHC*. These genes span transcriptional or chromatin regulation (*IRF2BPL, TFE3, FOXP3*, and *EZH2*) [17–20], intracellular signalling (*SH3BP2*) [21], and metabolism (*MMACHC*) [22]. Other genes, including *FH*, ranked highly for held-out gain but not for scale-adjusted CaLM weight. Given the limited variant counts within some genes, the rankings should be regarded as illustrative rather than as defining a fixed codon-dependent class.

Positive associations were also observed with ESM-1b (650M), both for mean CaLM weight and for CaLM AUROC after adjustment for PLM AUROC (Supplementary Figs. S8A and B). Similar patterns were obtained using the within-protein ESM-2 (150M) + ESM-2 (650M) ensemble as the reference (Supplementary Figs. S8C and D), supporting robustness to the protein-model background and the within-protein ensemble control.

These analyses indicate that the predictive benefit of adding CaLM varies across genes and remains associated with CaLM performance after adjustment for PLM performance. Within-gene cross-validation reduces optimism from weight selection, although limited per-gene sample sizes constrain the precision of individual estimates.

### 2.7 Matched DMS–CBGE comparisons suggest context-dependent shifts in inferred modality contribution

We next examined whether platform-dependent differences in CaLM contribution persisted after controlling for variant composition. DMS assays typically measure variant effects in exogenous or engineered expression systems, whereas CBGE assays evaluate variants at or near their endogenous genomic loci [16]. We identified 27 matched DMS–CBGE dataset pairs across 11 genes in which the same missense variants were evaluated on both platforms (Supplementary Table S6). Within each training fold, CaLM weights were optimised separately against the DMS and CBGE labels, scale-adjusted using the shared variants, and averaged across folds. Analyses were performed with ESM-2 (150M) and ESM-2 (650M) as alternative protein-model backgrounds (“Methods - Cross-platform controlled comparison”).

With ESM-2 (150M), 21 of 27 pairs showed higher CaLM weighting under CBGE, with a median Δ*w*^scale^ of 0.154 (95% gene-cluster bootstrap CI, 0.026–0.392; exact gene-block sign-flip *p* = 0.0156; Fig. 7A). With ESM-2 (650M), 18 pairs showed the same direction, although the median shift of 0.131 was less robust (95% CI, −0.017– 0.363; *p* = 0.0781; Fig. 7B).

**Fig. 7.**
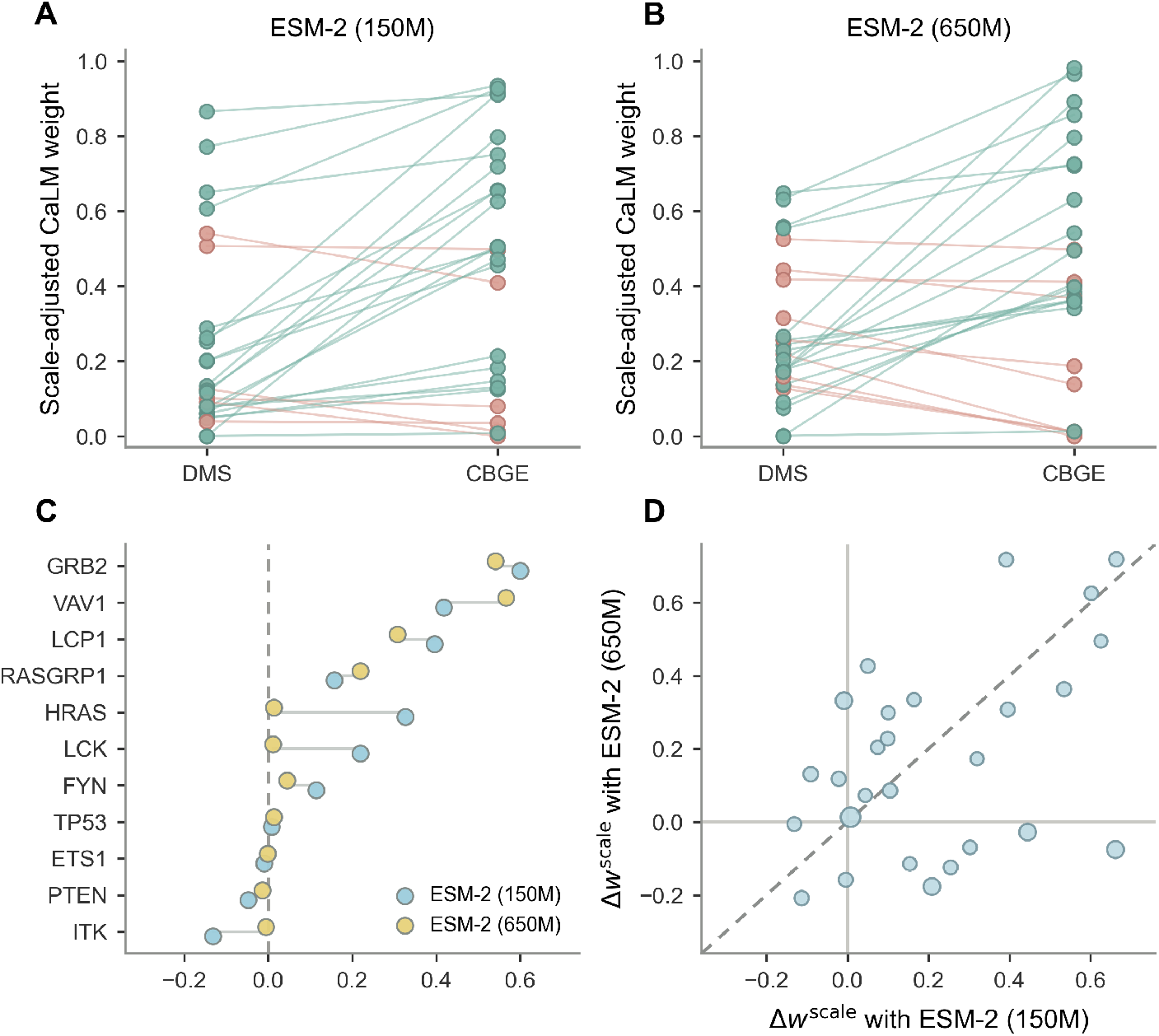
Matched ClinMAVE DMS–CBGE comparisons suggest context-dependent shifts in inferred modality contribution. **(A–B)** Paired comparison of scale-adjusted CaLM weights between matched DMS– CBGE dataset pairs using (**A**) ESM-2 (150M) or (**B**) ESM-2 (650M) as the PLM background. Each connected line represents one matched dataset pair. Green lines indicate a higher inferred CaLM weight under CBGE-derived labels, whereas coral lines indicate a lower weight. **(C)** Gene-level mean shift in scale-adjusted CaLM weight between CBGE and DMS, averaged across matched dataset pairs within each gene. Blue and pale-yellow points represent estimates obtained with ESM-2 (150M) and ESM-2 (650M), respectively; connecting lines link estimates for the same gene. **(D)** Concordance of pair-level Δ*w*^scale^ estimates between the two PLM backgrounds. Each point represents one matched dataset pair, with point size proportional to the number of shared variants. The dashed diagonal denotes equal shifts between backgrounds, while the horizontal and vertical reference lines indicate Δ*w*^scale^ = 0.

After averaging the two background-specific estimates within each pair, 24 of 27 pairs showed a positive shift. The median combined-background shift was 0.161 (95% CI, 0.043–0.293; *p* = 0.0313), supporting an overall tendency towards greater inferred CaLM contribution under CBGE-derived labels. However, shifts varied across genes and dataset pairs: *GRB2, VAV1*, and *LCP1* showed positive mean shifts under both backgrounds, whereas pairs from *HRAS* and *LCK* showed opposing directions (Fig. 7C; Supplementary Table S6).

Pair-level shifts were only moderately concordant between PLM backgrounds (Pearson *r* = 0.46; Fig. 7D). Thus, matched comparisons suggest a context-dependent increase in CaLM contribution under CBGE-derived labels, but not a universal platform effect.

## 3 Discussion

We asked whether CaLM-derived codon-level scores provide information beyond the residue-level representations of protein language models (PLMs) [1, 3–5]. In ClinVar, adding CaLM produced small but reproducible gains across several PLM backgrounds. These gains persisted after combining distinct PLMs and after controlling for explicit mutational-context features, although their magnitude decreased as the protein baseline became stronger. CaLM therefore provides a separable but limited source of predictive information rather than a substitute for protein-level scoring.

Same-modality PLM controls indicated that the CaLM gain was not solely attributable to the generic benefit of adding another predictor. Against the ESM-2 (650M) baseline, adding CaLM produced a larger gain than adding ESM-2 (150M), and a gain comparable to that obtained by adding ESM-2 (650M) to ESM-1b (650M), despite CaLM having substantially weaker performance. The distinction was smaller when ESM-1b itself served as the baseline, showing that the apparent benefit of CaLM depends on baseline strength. Its lower score correlation with ESM-2 is nevertheless consistent with CaLM capturing a partially distinct component of the predictive signal.

Mutational-context and probability-aggregation controls further clarified this contribution. CaLM and explicit codon-substitution features provided partially overlapping information, but CaLM retained a positive conditional contribution after these features were included. Aggregating CaLM probabilities across synonymous codons preserved the principal structure of CaLM–PLM disagreement while weakening its association with codon degeneracy. Synonymous probability splitting therefore contributes to substitution-level discordance, but neither this effect nor the low-order sequence features explains the broader ClinVar performance gain.

Across ClinMAVE, the inferred CaLM contribution depended on functional class, assay platform, gene, and PLM background. LoF prediction was largely protein-dominated, consistent with many LoF variants disrupting protein abundance, stability, folding, or molecular function [23–25]. Some GoF comparisons assigned high weight to CaLM, but these estimates were unstable and did not yield consistent held-out improvements. Matched DMS–CBGE comparisons showed a tendency towards higher CaLM weights under CBGE, although the direction and magnitude of the shift varied across dataset pairs and PLM backgrounds [16]. Optimised weights should therefore be interpreted as context-sensitive predictive parameters rather than fixed biological proportions.

Gene-level analyses supported the same interpretation. The benefit of adding CaLM varied continuously across genes and remained positively associated with CaLM performance after adjustment for PLM performance. This relationship was reproducible across ESM-2 and ESM-1b backgrounds and relative to a within-protein ensemble control. However, the identities and ordering of the highest-ranked genes were less stable across PLM backgrounds and analytical choices. These results identify gene-level contexts in which coding-sequence scores provide information not fully represented at amino-acid resolution, rather than a shared regulatory mechanism.

The nonsense–synonymous analysis provided a face-validity check for codon-level scoring. CaLM recovered the expected broad separation between nonsense and synonymous variants, consistent with purifying selection against premature termination and the relative tolerance of many synonymous substitutions [12, 26]. However, functional variation within either class was resolved only weakly. Such effects may depend on splicing, RNA structure, transcript stability, and cellular regulation, which are not directly represented by the coding-sequence scores evaluated here [27, 28].

This study focused on whether codon-level scores capture information complementary to protein-level representations. CaLM and the PLMs differ in scale, architecture, and training corpus, preventing attribution of the observed gains to codon resolution alone. The ClinMAVE analyses were further limited by sparse GoF classes and small gene-level sample sizes, and language-model scores do not establish causal mechanisms. Overall, CaLM-derived codon scores provide modest complementary information beyond protein-level representations. Matched-scale models and larger functional datasets will be needed to establish the generalisability and biological basis of this signal.

## 4 Methods

### 4.1 ClinVar variant dataset

ClinVar is a publicly available archive of clinically interpreted genetic variants and their supporting evidence [15]. Variant annotations were obtained from `variant summary.txt`, downloaded from the ClinVar FTP site on 8 October 2024 (https://ftp.ncbi.nlm.nih.gov/pub/clinvar). We retained single-nucleotide substitutions annotated as missense, nonsense, or synonymous variants with a clinical significance of *benign, likely benign, pathogenic*, or *likely pathogenic*, and a Clin-Var review status of at least one star. Variants with conflicting clinical-significance classifications were excluded.

RefSeq transcript identifiers and gene symbols were extracted from the standard variant descriptions in the ClinVar `Name` field. For genes with multiple transcripts, we retained the transcript containing the largest number of eligible variants and removed duplicate records. *Pathogenic* and *likely pathogenic* variants were grouped as pathogenic, whereas *benign* and *likely benign* variants were grouped as benign.

Because the primary model-combination analyses included ESM-1b, which accepts protein sequences of at most 1,022 residues, missense variants were restricted to proteins within this length range. We then retained complete cases with valid variant-level scores from CaLM, ESM-2 (150M), ESM-2 (650M), and ESM-1b (650M). The resulting matched missense cohort comprised 71,436 unique variants, including 24,933 pathogenic and 46,503 benign variants across 11,554 genes. This common cohort was used for the ClinVar model-combination, mutational-context, substitution-discordance, and gene-level analyses.

The nonsense and synonymous analyses did not require PLM compatibility because these variants were evaluated using CaLM alone. The corresponding cohorts comprised 46,386 nonsense variants across 3,291 genes and 527,580 synonymous variants across 11,593 genes. Cohort composition for each analysis-specific dataset is summarised in Fig. 1C and Supplementary Table S1.

### 4.2 ClinMAVE variant dataset

ClinMAVE is a curated resource that harmonises functional evidence from multiplexed assays of variant effect and assigns variants to functionally normal, LoF, or GoF categories using reviewed assay-specific thresholds [16]. ClinMAVE records were accessed on 15 March 2026. We collected missense, nonsense, and synonymous variants and grouped them by experimental platform into DMS and CBGE datasets.

For missense analyses involving ESM-1b (650M), variants were restricted to proteins of no more than 1,022 residues. Duplicate records were collapsed by variant identifier within each platform and functional class. For functional-class comparisons, variants assigned conflicting functional classes across ClinMAVE source files were excluded before constructing the functionally normal versus LoF and functionally normal versus GoF datasets.

The resulting missense dataset comprised 168,089 DMS variants across 311 genes and 90,242 CBGE variants across 309 genes. The CaLM-only analyses included 11,021 DMS and 6,475 CBGE nonsense variants across 418 and 399 genes, respectively, together with 12,666 DMS and 45,916 CBGE synonymous variants across 23 and 399 genes, respectively. Counts by functional category and platform are provided in Figs. 1D and E and Supplementary Table S2.

### 4.3 CaLM and PLMs

To compare codon- and protein-level representations, we employed CaLM (86 million parameters; downloaded from https://github.com/oxpig/CaLM) [13] together with PLMs from the ESM family. We evaluated ESM-2 at two scales: ESM-2 (150 million parameters; downloaded from https://huggingface.co/facebook/esm2_t30_150M_UR50D) and ESM-2 (650 million parameters; downloaded from https://huggingface.co/facebook/esm2_t33_650M_UR50D) [3]. For model-control analyses, we additionally evaluated ESM-1b (650 million parameters; downloaded from https://huggingface.co/facebook/esm1b_t33_650M_UR50S) [5]. ESM-1b provided a protein-level comparator with a different architecture and training corpus from ESM-2. These models differ in three key respects:

1. Tokens: CaLM tokenises cDNA sequences into codons, whereas ESM models tokenise protein sequences into amino-acid residues.
2. Training data: CaLM was trained on cDNA sequences from the European Nucleotide Archive, whereas ESM models were trained on large protein-sequence databases derived from UniRef.
3. Output logits: CaLM generates position-specific logits over codon tokens, whereas ESM models generate position-specific logits over amino-acid tokens.

### 4.4 Metrics for variant classification

For ClinVar classification, pathogenic variants were treated as the positive class and benign variants as the negative class. For ClinMAVE classification, functionally normal variants were treated as the negative class, while LoF and GoF variants were separately treated as the positive class in the corresponding binary tasks. Classification performance was evaluated using AUROC.

### 4.5 Missense effect scores

Following Brandes et al. [4], we used a single-pass wt-marginal scoring strategy. The unmasked wt coding sequence was provided to CaLM, and the corresponding wt protein sequence was provided to each PLM. At every sequence position, each model produced logits over its respective codon or amino-acid vocabulary, which were converted to probabilities using softmax. This procedure allowed the LLRs of all possible single-token substitutions to be calculated from one forward pass per sequence.

For a missense variant at position *k*, let 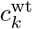 and 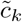 denote the wt and mutant codons, respectively, and let 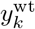 and 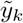 denote the corresponding wt and mutant amino acids. The CaLM codon-level LLR was defined as

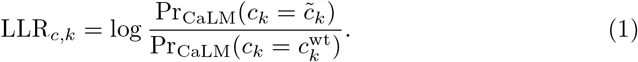

For PLM *m*, the residue-level LLR was defined analogously as

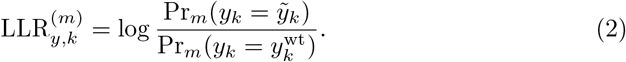

Because lower LLR values indicate that the mutant token is less probable than the wt token, pathogenicity-oriented effect scores were defined as

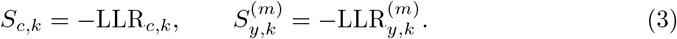

These scores were used directly for single-model analyses. Model complementarity was evaluated using convex linear ensembles:

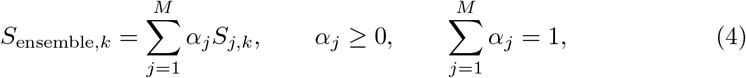

where *S*_*j,k*_ denotes a CaLMor PLM-derived pathogenicity score and *M* denotes the number of component models. Scores were combined on their original scales.

Because raw ensemble weights depend on the dispersion of the component scores, we then reported scale-adjusted weights for descriptive comparisons. For component *q* in training fold *f*, let *α*_*q,f*_ denote its raw weight and *σ*_*q,f*_ denote the standard deviation of its training-set scores. The scale-adjusted weight was defined as

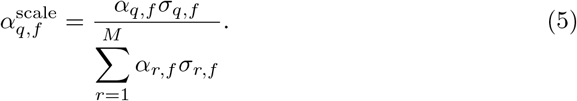

These values represent the relative component coefficients when the same fitted ensemble is expressed in terms of training-set z-scored component scores. Mean centring introduces an additive constant that does not affect variant ranking or AUROC. Scale-adjusted weights were used only for reporting and cross-context comparisons; all out-of-fold predictions and AUROC calculations used the original raw-score ensemble weights.

### 4.6 Model evaluation and ensemble optimisation

Unless otherwise specified, performance was evaluated using 10-fold gene-held-out cross-validation with `StratifiedGroupKFold` in `scikit-learn` (v1.6.1) [29]. All variants from the same gene were assigned to the same fold, and identical splits were used across models. Held-out predictions were pooled across folds and AUROC was calculated from the combined out-of-fold predictions. Folds containing only one class were excluded from fold-level analyses but retained in pooled AUROC calculations. Analysis-specific cross-validation schemes are described separately below.

Convex ensemble weights were optimised independently within each training split by grid search to maximise AUROC and then applied unchanged to the corresponding held-out data. Two-model ensembles used weights from 0 to 1 in increments of 0.01, whereas three-model ensembles were searched over the probability simplex in increments of 0.05.

CIs for ΔAUROC were estimated from 5,000 gene-cluster bootstrap replicates of the pooled out-of-fold predictions. Genes were sampled with replacement, retaining all variants within each sampled gene, and paired AUROC differences were recalculated for each replicate without re-optimising ensemble weights. Percentile-based 95% CIs were reported. This procedure preserved within-gene dependence and avoided treating overlapping cross-validation folds as independent.

To distinguish cross-modal complementarity from the generic benefit of ensembling, PLM-only controls were evaluated using the same cross-validation and optimisation procedure. ESM-2 (150M) was combined with ESM-2 (650M), and ESM-2 (650M) with ESM-1b (650M), with gains measured relative to the stronger constituent model. CaLM gain was defined analogously relative to ESM-2 (650M). All gains were calculated from pooled out-of-fold predictions, with uncertainty estimated by genecluster bootstrap. Score redundancy between each added scorer and ESM-2 (650M) was assessed using Spearman rank correlation across the ClinVar missense dataset (Supplementary Table S4).

### 4.7 Codon-level mutational-context control analysis

We derived mutational-context covariates from the reference and mutant codons of each ClinVar missense variant. Codons were converted to the DNA alphabet where necessary by replacing U with T. We retained only valid single-nucleotide codon substitutions for which both codons were three nucleotides long, contained only A, C, G, and T, and differed at exactly one position.

For each variant, categorical covariates included the reference and mutant codons, the mutated codon position, the reference and mutant nucleotides, and the nucleotide-substitution class (e.g., A → G). Additional covariates included transition/transversion status, reference and mutant codon GC content, the change in codon GC content, presence of a CpG dinucleotide in the reference and mutant codons, the change in within-codon CpG status, and local codon degeneracy. Local codon degeneracy was defined as the number of nucleotide identities, including the reference nucleotide, that could occur at the mutated codon position without changing the amino acid encoded by the reference codon under the standard genetic code.

Mutational-context controls were evaluated using the same 10-fold gene-held-out cross-validation as the model-combination analyses. Within each training split, numeric covariates were standardised and categorical covariates were one-hot encoded, and the fitted transformations were then applied to the corresponding held-out genes. ESM-2 (650M) and CaLM were represented by their respective pathogenicity scores. Using an L2-regularised logistic-regression classifier, we compared four feature sets: ESM-2 (650M) alone, ESM-2 (650M) with mutational-context covariates, ESM-2 (650M) with CaLM, and ESM-2 (650M) with both mutational-context covariates and CaLM. Conditional contributions were quantified as differences in AUROC calculated from pooled out-of-fold predictions, with 95% CIs estimated by gene-cluster bootstrap.

### 4.8 Characterisation of cross-modal substitution discordance

To compare CaLM and ESM-2 (650M) in a common amino-acid probability space, CaLM codon probabilities were summed over synonymous codons. For position *k*, the aggregated probability and corresponding LLR were defined as

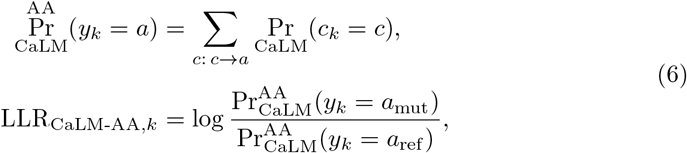

where *c* → *a* denotes codons encoding amino acid *a*, and *a*_ref_ and *a*_mut_ denote the reference and mutant amino acids, respectively.

Cross-modal discordance was defined as

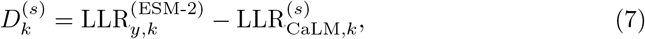

where *s* denotes codon or amino-acid-aggregated space, and 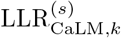 denotes LLR_*c,k*_ or LLR_CaLM-AA,*k*_, respectively. Positive values indicate a higher LLR under ESM-2 than under CaLM.

Variants with discordance scores below the 1st percentile or above the 99th percentile in each space were classified as extremely discordant. Pathogenic variants in the upper tail were designated CaLM-leaning, whereas those in the lower tail were designated PLM-leaning; these assignments were reversed for benign variants.

The association between codon degeneracy and CaLM-leaning discordance was evaluated separately in each probability space using logistic regression:

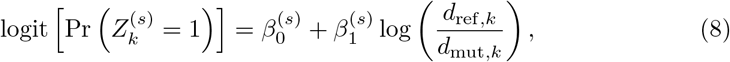

where 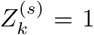 identifies CaLM-leaning extreme variants and 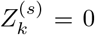 identifies all other scored variants. The quantities *d*_ref,*k*_ and *d*_mut,*k*_ denote the numbers of codons encoding the reference and mutant amino acids, respectively. ORs were calculated as 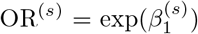. Standard errors were clustered by gene to account for within-gene dependence, and 95% CIs for the ORs were obtained by exponentiating the corresponding coefficient confidence limits. An OR *<* 1 indicates lower odds of CaLM-leaning discordance per one-unit increase in the log reference-to-mutant degeneracy ratio.

Substitution-level enrichment within each discordant subset *S*_*i*_ was quantified as a background-normalised ratio (BR),

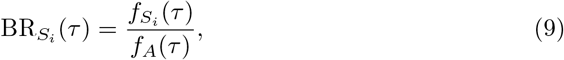

where 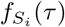 and *f*_*A*_(*τ*) denote the frequencies of reference-to-mutant amino acid substitution *τ* in *S*_*i*_ and in the complete scored ClinVar missense cohort *A*, respectively. For each substitution, a two-sided Fisher’s exact test compared its occurrence in *S*_*i*_ with that in *A \ S*_*i*_. Benjamini–Hochberg correction was applied across tested substitution pairs separately within each discordant subset. Substitutions with adjusted *p <* 0.01 and 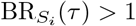 were considered significantly enriched.

### 4.9 Gene-level codon contribution analysis

Gene-level analyses included 466 genes with at least five pathogenic and five benign variants in the ClinVar missense cohort. For each gene *g*, CaLM and PLM scores were combined as

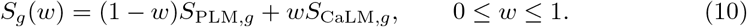

Performance was evaluated using stratified five-fold cross-validation within each gene. For each fold *f*, the CaLM weight was selected by maximising AUROC on the remaining four folds:

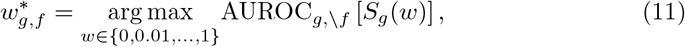

where *g, \ f* denotes the training variants for gene *g*. The selected weight was applied without refitting to the held-out fold. Out-of-fold predictions were pooled across all five folds to calculate the cross-validated ensemble AUROC, AUROC^CV^. The selected raw weight was converted to its scale-adjusted value using the two score standard deviations in the corresponding training split. The mean scale-adjusted weight across folds was denoted 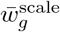.

With ESM-2 (650M) as the primary PLM background, cross-modal gain was defined as

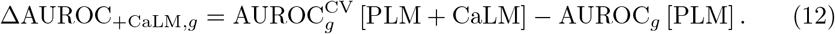

Both AUROCs were calculated over the same variants, using out-of-fold ensemble predictions and the corresponding fixed PLM scores.

The association between 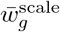 and cross-modal gain was evaluated using ordinary least-squares regression. We also repeated the regression using weighted least squares, giving greater weight to genes with larger and more balanced variant sets:

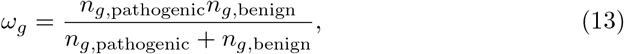

where the two counts denote the numbers of pathogenic and benign variants in gene *g*. Both analyses used heteroskedasticity-consistent HC3 estimates for standard errors, 95% CIs, and *p*-values.

To assess the association between CaLM performance and cross-modal gain conditional on PLM performance, we fitted

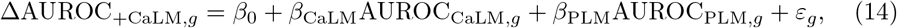

where AUROC_CaLM,*g*_ and AUROC_PLM,*g*_ denote the individual-model AUROCs for gene *g*, and *β*_CaLM_ quantifies the association after adjustment for PLM AUROC.

For visualisation, CaLM AUROC and cross-modal gain were separately regressed on PLM AUROC, including an intercept. The resulting residuals, *r*_CaLM,*g*_ and *r*_gain,*g*_, were plotted against one another to display the adjusted association.

Analyses were repeated using ESM-1b (650M) as an alternative PLM background. As a within-protein ensemble control, we additionally evaluated the cross-validated AUROC difference between ESM-2 (650M) + CaLM and ESM-2 (150M) + ESM-2 (650M).

### 4.10 Cross-platform controlled comparison

Matched DMS–CBGE datasets were obtained from ClinMAVE, accessed on 15 March 2026 [16]. We paired experiments assaying missense variants in the same gene and restricted the analysis to functionally normal and LoF variants. Candidate pairs required at least five variants per class on each platform. Within each pair, we retained shared variants with unambiguous labels and valid CaLM and ESM-2 scores, yielding 27 pairs across 11 genes.

Cross-validation was performed within each pair using shuffled `StratifiedKFold` with a fixed seed of 16. Variants were stratified by their joint DMS–CBGE labels, and the number of folds was the smaller of 10 and the size of the smallest observed stratum. Pairs lacking both classes on either platform or supporting fewer than two folds were excluded. Fold assignments were shared across platforms and PLM backgrounds.

Within each training fold, CaLM weights were optimised separately against DMS and CBGE labels by grid search in increments of 0.005 and scale-adjusted using the shared training-score standard deviations. The platform-associated shift for pair *p* was defined as

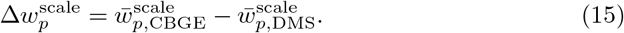

Positive values indicate greater CaLM weighting under CBGE-derived labels. A cross-background summary was obtained by averaging the shifts from ESM-2 (150M) and ESM-2 (650M):

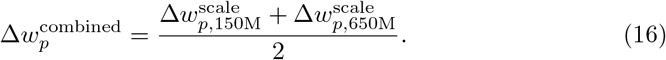

The median combined-background shift and its 95% CI were estimated using 10,000 gene-cluster bootstrap replicates. Significance was assessed using a two-sided exact gene-block sign-flip test with the median shift as the test statistic and a common sign assigned to all pairs from the same gene. The same procedures were applied to the background-specific estimates as sensitivity analyses, while their pair-level correlation was summarised descriptively using Pearson’s *r*.

### 4.11 Data analysis and visualisation

Data analysis and figure preparation were performed using Python v3.9.13. The main Python libraries used were NumPy v1.24.4, pandas v2.3.3, SciPy v1.9.1, scikit-learn v1.6.1, statsmodels v0.13.2, PyTorch v2.2.2, Matplotlib v3.9.4, and seaborn v0.13.2.

### 4.12 Use of large language models

During the preparation of this manuscript, the authors used Claude Opus 4.8 (Anthropic) to improve the clarity, grammar, and readability of the text.

## Acknowledgements

Figure 1 was created with BioRender (https://www.biorender.com/).

## Author contributions

R.C. and M.B. designed the study. R.C. implemented code and methods. M.B., G.F. and N.P. supervised the project. All authors contributed to the writing, reviewed, and approved the final manuscript.

## Conflict of interest

None declared.

## Funding

This work was supported by Australian Government Research Training Program Scholarship to R.C. N.P. and M.B. received support from Medical Research Future Fund (2016033).

## Data availability

All the data and code necessary to reproduce the analyses, figures, and results are available on our GitHub repository: https://github.com/Cassie818/Viral-mut.

## Notes

### Competing Interest Statement

The authors have declared no competing interest.

### Summary of Updates

This version has been revised to strengthen the robustness of the analyses and clarify the main conclusions. We added gene level bootstrap confidence intervals, revised the gene level analysis to control for baseline PLM performance, and included additional controls for mutational context, score scaling, and PLM ensembling.

