## Supplementary material for "Codon language model scores provide information beyond protein language models for missense variant interpretation": supp

#### Supplementary Tables

| | **Missense mutation** | **Nonsense mutation** | **Synonymous mutation** |
|---|---|---|---|
| Pathogenic | 9,614 | 37,113 | 213 |
| Likely pathogenic | 15,319 | 8,834 | 219 |
| Benign | 14,208 | 134 | 38,903 |
| Likely benign | 32,295 | 305 | 488,245 |
| Total | 71,436 | 46,386 | 527,580 |
| Genes | 11,554 | 3,291 | 11,593 |

**Supplementary Table S1** Three types of single-point mutations collected from ClinVar.

| MAVE technique | Missense DMS | Missense CBGE | Nonsense DMS | Nonsense CBGE | Synonymous DMS | Synonymous CBGE |
|---|---|---|---|---|---|---|
| Functionally normal | 117,412 (311) | 82,522 (273) | 697 (73) | 5,234 (396) | 11,983 (23) | 32,467 (172) |
| LoF | 49,258 (309) | 5,650 (112) | 10,276 (416) | 1,198 (59) | 454 (13) | 7,286 (268) |
| GoF | 1,419 (8) | 2,070 (111) | 48 (1) | 43 (14) | 229 (3) | 6,163 (265) |
| Total | 168,089 (311) | 90,242 (309) | 11,021 (418) | 6,475 (399) | 12,666 (23) | 45,916 (399) |

**Supplementary Table S2** Three types of single-point mutations collected from ClinMAVE.
Values in parentheses indicate the number of genes.

| | Pathogenic vs. Benign | | Functionally normal vs. LoF | | | | Functionally normal vs. GoF | | | |
|---|---|---|---|---|---|---|---|---|---|---|
| | | | DMS | | CBGE | | DMS | | CBGE | |
| | Mean ($\mu$) | Std dev ($\sigma$) | Mean ($\mu$) | Std dev ($\sigma$) | Mean ($\mu$) | Std dev ($\sigma$) | Mean ($\mu$) | Std dev ($\sigma$) | Mean ($\mu$) | Std dev ($\sigma$) |
| CaLM | -5.37 | 2.83 | -6.28 | 2.64 | -5.33 | 2.09 | -5.78 | 2.39 | -5.24 | 2.00 |
| ESM-2 (650M) | -6.21 | 3.63 | -7.37 | 3.35 | -5.94 | 3.09 | -6.57 | 3.11 | -5.75 | 2.95 |
| ESM-1b (650M) | -7.25 | 4.56 | -9.59 | 4.24 | -7.57 | 4.01 | -8.72 | 4.12 | -7.51 | 3.96 |
| $\sigma$ (CaLM) / $\sigma$ (ESM-2) | 0.78 | | 0.79 | | 0.68 | | 0.77 | | 0.68 | |
| $\sigma$ (CaLM) / $\sigma$ (ESM-1b) | 0.62 | | 0.62 | | 0.52 | | 0.58 | | 0.50 | |

**Supplementary Table S3** Distributional scale of model LLR outputs used for ensemble analyses.

| Added scorer | Role | Spearman rho with ESM-2 (650M) | Pearson r |
| --- | --- | --- | --- |
| ESM-2 (150M) | weak PLM placeholder | 0.8365 | 0.8433 |
| ESM-1b (650M) | strong PLM placeholder | 0.8300 | 0.8208 |
| CaLM | cross-modal codon scorer | 0.6312 | 0.6793 |

**Supplementary Table S4** Score correlations with the ESM-2 (650M) baseline. Spearman and Pearson correlations between each added scorer and ESM-2 (650M) scores across 71,436 ClinVar missense variants. ESM-2 (150M) and ESM-1b (650M) serve as protein-modality placeholder controls, whereas CaLM represents the codon-level scorer.

| Covariate | Encoding | Definition |
| --- | --- | --- |
| Reference codon | Categorical | Wild-type DNA codon (U converted to T). |
| Mutant codon | Categorical | Mutant DNA codon (U converted to T). |
| Mutated codon position | Categorical | Position 1, 2, or 3 within the codon. |
| Reference nucleotide | Categorical | Wild-type nucleotide at the mutated position. |
| Mutant nucleotide | Categorical | Mutant nucleotide at the mutated position. |
| Nucleotide substitution | Categorical | Directed substitution, for example C>T. |
| Transition status | Numeric | 1 for A>G, G>A, C>T, or T>C; otherwise 0. |
| Transversion status | Numeric | 1 minus transition status. |
| Reference codon GC content | Numeric | Fraction of G or C nucleotides in the reference codon. |
| Mutant codon GC content | Numeric | Fraction of G or C nucleotides in the mutant codon. |
| Change in codon GC content | Numeric | Mutant minus reference codon GC content. |
| Reference within-codon CpG status | Numeric | 1 if the reference codon contains CG; otherwise 0. |
| Mutant within-codon CpG status | Numeric | 1 if the mutant codon contains CG; otherwise 0. |
| Change in within-codon CpG status | Numeric | Mutant minus reference within-codon CpG status. |
| Local codon degeneracy | Categorical | Number of possible nucleotides at the mutated position that preserve the reference amino acid. |

**Supplementary Table S5** Mutational-context covariates used in the context-control analysis.
Numeric covariates were standardised and categorical covariates were one-hot encoded using training genes within each cross-validation fold.

| Gene | DMS dataset | CBGE dataset | N | Label concordance | Δw (150M) | Δw (650M) |
|---|---|---|---|---|---|---|
| ETS1 | dataset0541 | dataset0537 | 59 | 0.627 | 0.105 | 0.086 |
| ETS1 | dataset0541 | dataset0538 | 59 | 0.695 | -0.021 | 0.118 |
| ETS1 | dataset0541 | dataset0539 | 58 | 0.638 | -0.113 | -0.208 |
| FYN | dataset0631 | dataset0629 | 33 | 0.636 | 0.075 | 0.204 |
| FYN | dataset0631 | dataset0627 | 32 | 0.688 | 0.154 | -0.115 |
| GRB2 | dataset0654 | dataset0650 | 59 | 0.729 | 0.664 | 0.718 |
| GRB2 | dataset0654 | dataset0652 | 54 | 0.648 | 0.535 | 0.363 |
| HRAS | dataset0688 | dataset0691 | 148 | 0.486 | 0.662 | -0.076 |
| HRAS | dataset0685 | dataset0691 | 140 | 0.657 | 0.208 | -0.177 |
| HRAS | dataset0686 | dataset0691 | 138 | 0.406 | -0.009 | 0.331 |
| HRAS | dataset0687 | dataset0691 | 138 | 0.536 | 0.445 | -0.028 |
| ITK | dataset0812 | dataset0810 | 45 | 0.622 | -0.131 | -0.006 |
| LCK | dataset0914 | dataset0913 | 45 | 0.533 | 0.255 | -0.124 |
| LCK | dataset0914 | dataset0912 | 43 | 0.581 | 0.099 | 0.228 |
| LCK | dataset0914 | dataset0910 | 37 | 0.514 | 0.302 | -0.070 |
| LCP1 | dataset0920 | dataset0918 | 48 | 0.75 | 0.396 | 0.307 |
| PTEN | dataset1477 | dataset1481 | 63 | 0.651 | -0.091 | 0.131 |
| PTEN | dataset1476 | dataset1481 | 51 | 0.627 | -0.004 | -0.158 |
| RASGRP1 | dataset1562 | dataset1558 | 27 | 0.519 | 0.044 | 0.072 |
| RASGRP1 | dataset1562 | dataset1559 | 27 | 0.63 | 0.164 | 0.334 |
| RASGRP1 | dataset1562 | dataset1560 | 26 | 0.385 | 0.101 | 0.299 |
| RASGRP1 | dataset1562 | dataset1561 | 26 | 0.423 | 0.320 | 0.173 |
| TP53 | dataset1982 | dataset1983 | 1195 | 0.777 | 0.008 | 0.013 |
| VAV1 | dataset2065 | dataset2064 | 46 | 0.696 | 0.392 | 0.717 |
| VAV1 | dataset2065 | dataset2061 | 45 | 0.689 | 0.050 | 0.426 |
| VAV1 | dataset2065 | dataset2062 | 37 | 0.703 | 0.602 | 0.625 |

| | | | | | | |
|---|---|---|---|---|---|---|
| VAV1 | dataset2065 | dataset2063 | 20 | 0.75 | 0.625 | 0.494 |

**Supplementary Table S6** Matched DMS-CBGE dataset pairs used for experimental-context analysis.

### Supplementary Figures

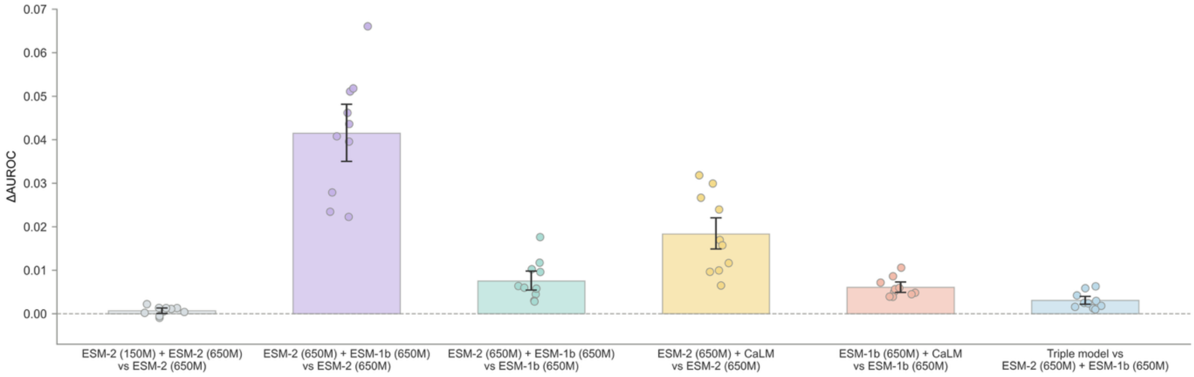


**Fig S1** AUROC gains and gene-cluster bootstrap uncertainty for model-combination analyses.

Coloured points show fold-level AUROC differences between each combined model and its corresponding baseline. Black points denote differences calculated from pooled out-of-fold predictions, and error bars indicate 95% percentile confidence intervals from 5,000 gene-cluster bootstrap replicates.

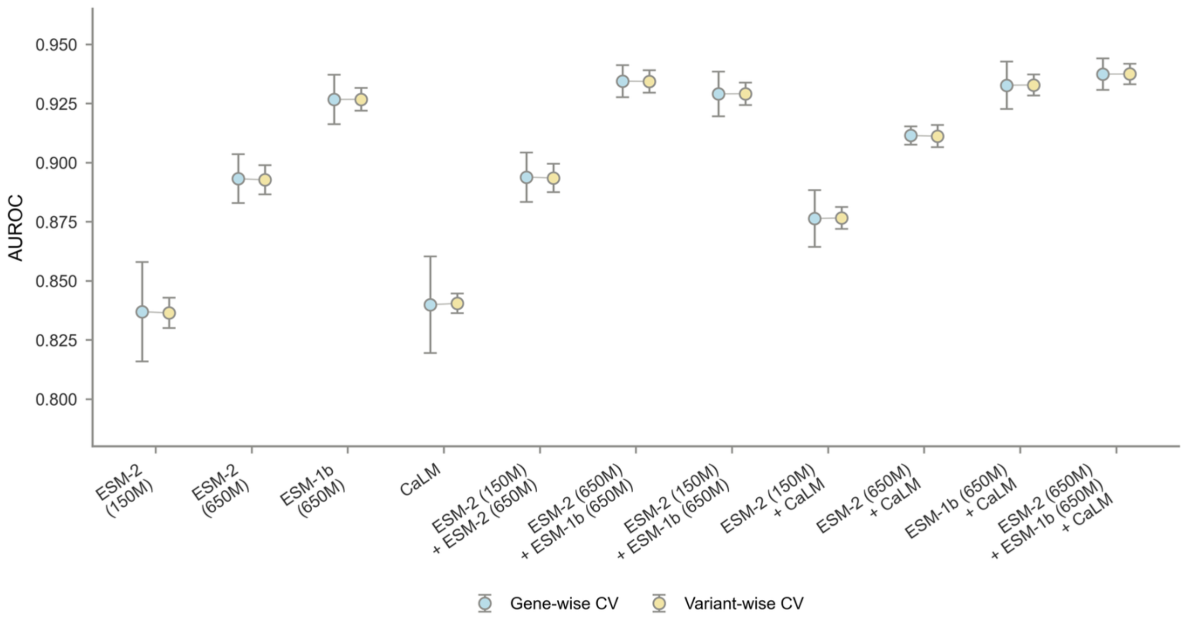


**Fig S2** Comparison of gene-wise and variant-wise cross-validation performance. Mean AUROC values are shown for each model under gene-wise cross-validation and variant-wise cross-validation. Error bars indicate the standard deviation across folds.

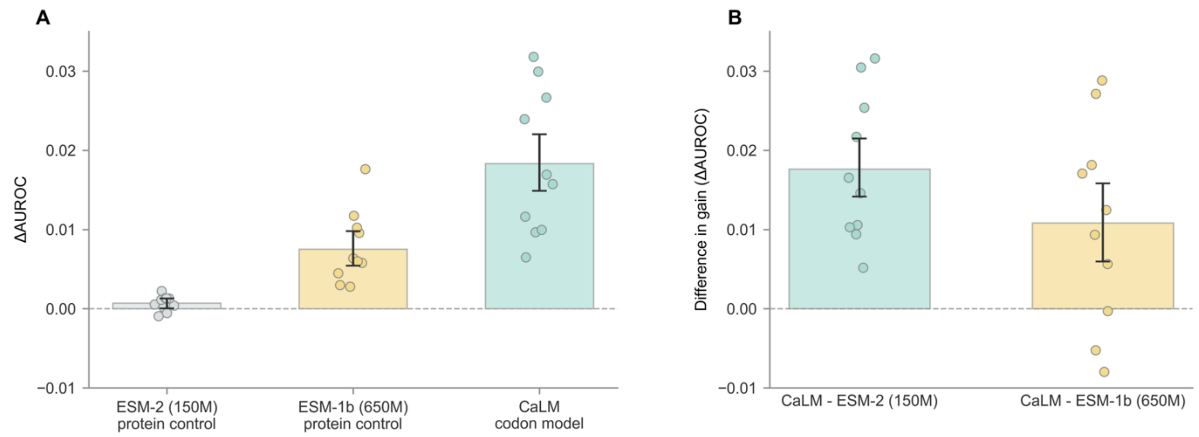


**Fig S3** Same-modality controls for generic model-ensembling effects.

**(A)** AUROC gains obtained by adding ESM-2 (150M) or CaLM to ESM-2 (650M), or by adding ESM-2 (650M) to the stronger ESM-1b (650M) baseline. Coloured points represent estimates from individual gene-held-out folds, grey lines connect estimates from the same fold across comparisons, and black points denote gains calculated from the pooled out-of-fold predictions.

**(B)** Differences between the CaLM gain and each same-modality PLM control gain. Coloured points show fold-level differences. Black points denote differences calculated from the pooled out-of-fold predictions, and horizontal error bars indicate 95% confidence intervals estimated from 5,000 gene-cluster bootstrap replicates.

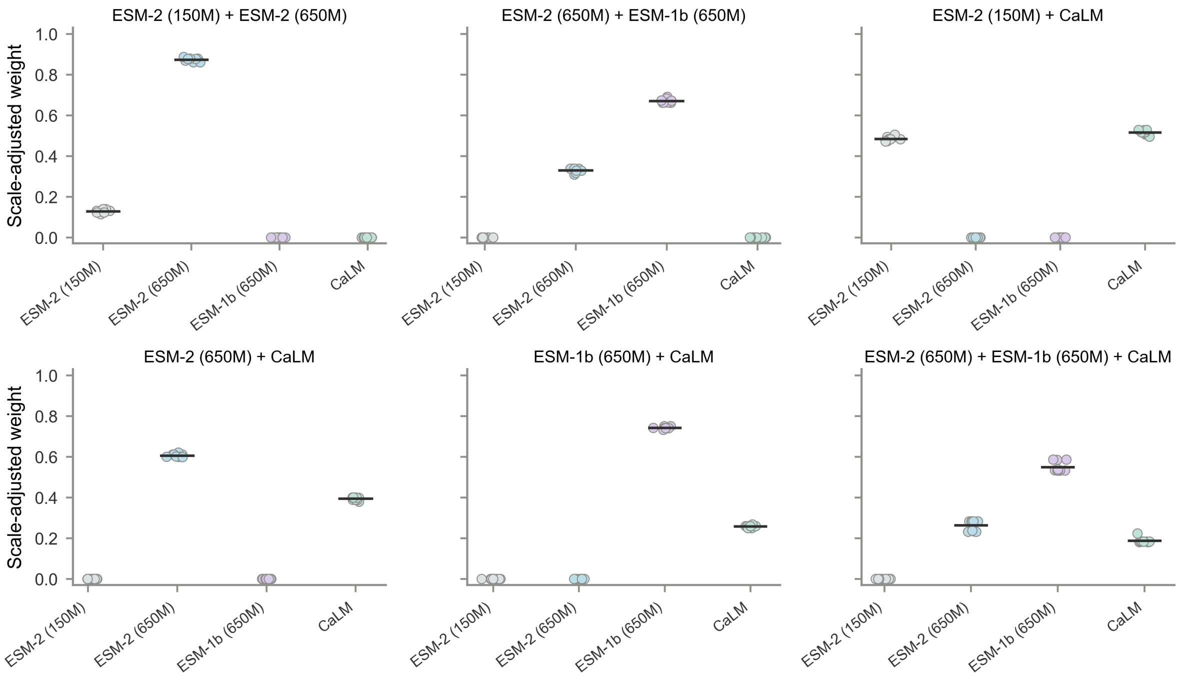


**Fig S4** Fold-level ensemble weights in model-combination analyses.
Optimised model weights are shown across the 10 gene-wise folds for each ensemble.

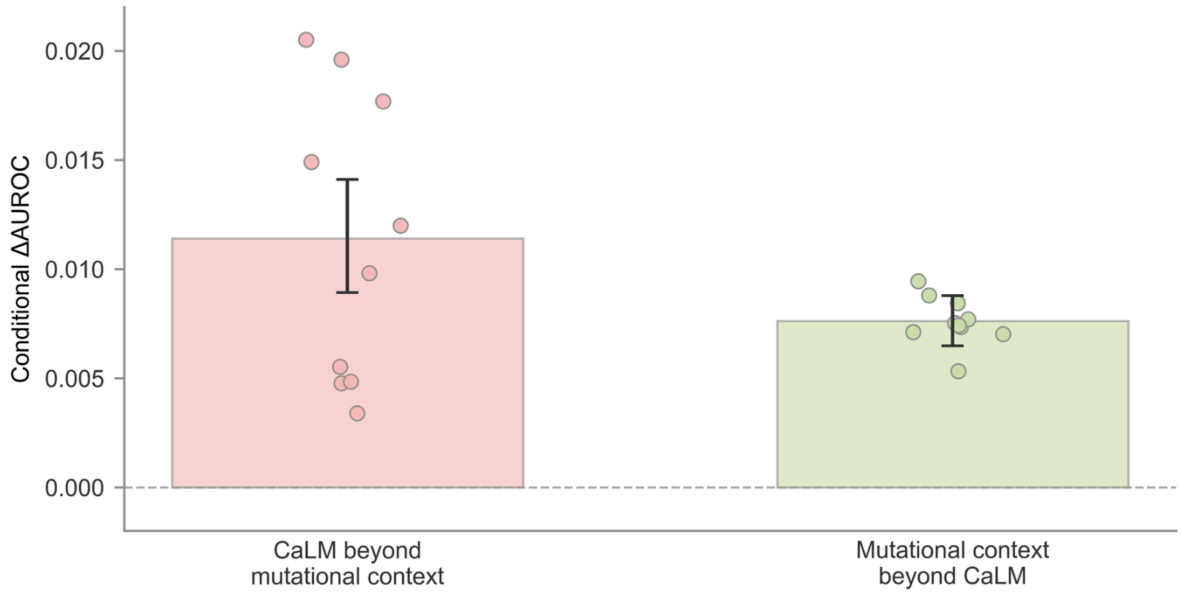


**Fig. S5** Conditional contributions of CaLM and explicit mutational context.

Coloured points represent conditional $\Delta AUROC$ values from individual gene-held-out folds. Black points denote gains calculated from pooled out-of-fold predictions, and horizontal error bars indicate 95% confidence intervals from 5,000 gene-cluster bootstrap replicates.

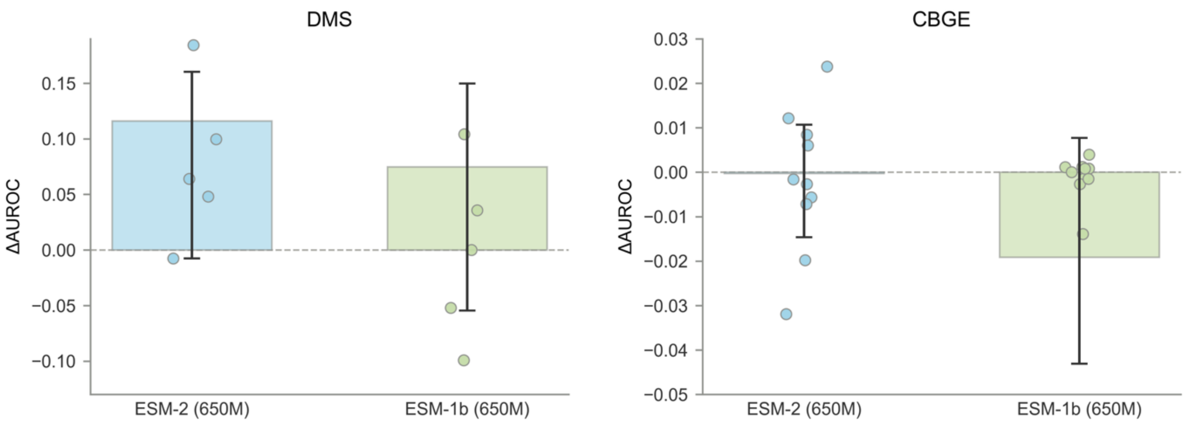


**Fig. S6** AUROC gains and uncertainty for ClinMAVE GoF comparisons.
Coloured points represent $\Delta AUROC$ values from evaluable gene-held-out folds. Black points denote gains calculated from pooled out-of-fold predictions, and horizontal error bars indicate 95% confidence intervals from 5,000 gene-cluster bootstrap replicates.

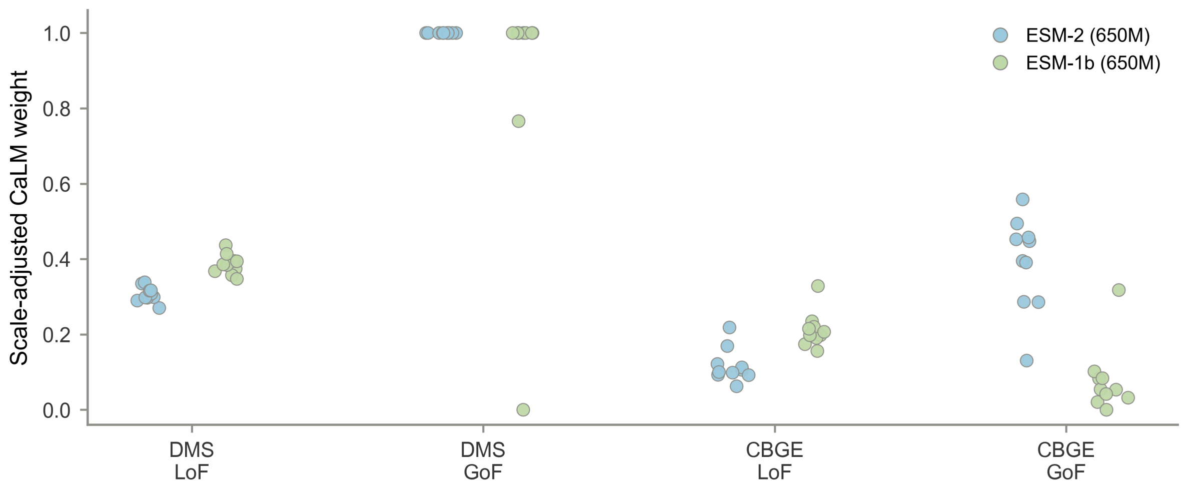


**Fig S7** Per-fold optimised CaLM weights in ClinMAVE functional-class analyses. Each point represents the CaLM weight selected using training genes in one gene-held-out cross-validation fold. Blue and green points indicate CaLM ensembles with ESM-2 (650M) and ESM-1b (650M), respectively.

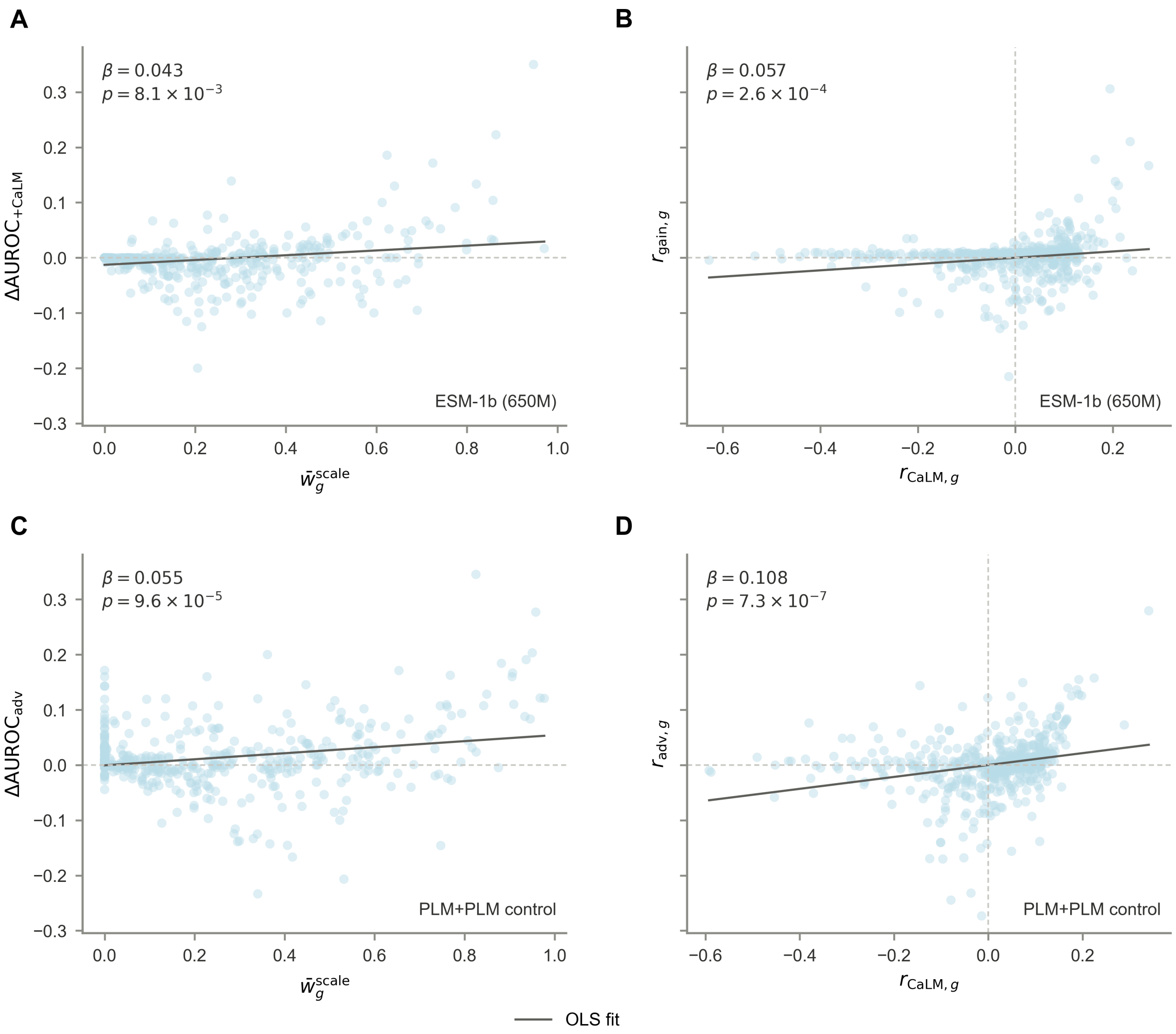


**Fig. S8** Gene-level CaLM contribution is robust to PLM-background and ensemble-control analyses.

**(A)** Association between mean CaLM weight and held-out AUROC gain using ESM-1b (650M) as the PLM background.

**(B)** Partial-regression association between CaLM AUROC and cross-modal gain after controlling for ESM-1b (650M) AUROC.

**(C)** Association between and cross-modal advantage, defined relative to the ESM-2 (150M) + ESM-2 (650M) ensemble.

**(D)** Partial-regression association between CaLM AUROC and cross-modal advantage after controlling for the AUROC of the PLM-only ensemble. Each point represents one gene, and solid lines indicate OLS fits.
